# Environmental conditions predetermine innate antioxidants pool in sea oat (*Uniola paniculata* L.) seeds

**DOI:** 10.1101/2023.04.11.536396

**Authors:** Andrew Ogolla Egesa, Héctor E. Pérez, Kevin Begcy

## Abstract

**Background and Aims:** Secondary metabolites such as antioxidants are critical components that protect seeds from stress damage during seed development, desiccation, and ex-situ storage. Antioxidants are essential determinants of seed quality, longevity, and persistence. Understanding the environmental factors that regulate the accumulation, content, and function of antioxidant pools in sea oat seeds is critical for gene banking and understanding the environmental impacts on seed quality.

**Methods:** Germination, viability, and Trolox Equivalent Antioxidant Capacity (TEAC) were analyzed on seeds from 18 sea oat populations from the US Atlantic and Gulf of Mexico coasts. We first assessed baseline TEAC, followed by TEAC performed on imbibed seeds for 36 hours at 35/25 °C during the day and night, respectively. Then, we analyzed the relationship of the antioxidant pools from these 18 sea oat populations to sampling site environmental classifications.

**Key Results:** Higher baseline antioxidants were common in regions with extreme environmental conditions corresponding to sea oat populations growing at latitudinal extremes characterized by warmer and colder temperatures. Baseline antioxidants did not correlate with prevailing seed germination. However, higher concentrations of antioxidants following imbibition were associated with poor seed germination in warmer conditions.

**Conclusions:** Our results indicate that climatic conditions and environmental components associated with temperatures and precipitation may largely influence the innate pool of antioxidants in sea oat seeds. Also, a high amount of antioxidants following sea oat seeds imbibition suggest seed damage or poor viability influenced by environmental stress conditions during seed development.

## INTRODUCTION

*Uniola paniculata* L. (sea oats [Poaceae]) is an ecological keystone species occurring in coastal dune systems throughout the Yucatan peninsula, Cuba, the Bahamas, and southeastern US coastlines (Subudhi et al. 2005; Lonard et al. 2011; Hodel and Gonzales 2013). Sea oats are essential for their ecological services, such as sand accretion, preventing erosion, and forming and stabilizing coastal dunes. These services largely buffer the effect of extreme events such as storms and hurricanes prevalent along those coastlines (Lonard et al. 2011; Hacker et al. 2019). Consequently, sea oats are in high demand for habitat restoration and re-vegetation programs. However, the spatial extent of sea oats populations has diminished greatly due to habitat destruction, conversion, and fragmentation. Protecting remaining sea oats populations is therefore crucial for maintaining donor populations for seed-based conservation and restoration programs.

Sea oats face diverse extreme conditions in the coastal dune habitat including osmotic stress from aerosolized salts and saline soils, hypoxia due to flooding, high and low-temperature stress, high humidity stress, and physical damage from tidal and wave over-wash or winds caused by storms and hurricanes. Sea oats also face burial from shifting dune sands. Therefore, sea oats and their seeds need highly functional stress-relief systems, especially antioxidants, to survive such conditions. However, there is limited knowledge on the presence, environmental influence, and functional role of antioxidants in plants from coastal dune systems exposed to such wide-ranging stressors.

Antioxidants are a category of secondary metabolites synthesized by living organisms that have been associated with preventing oxidative damage to cells and tissues. These molecules function at relatively low concentrations during optimal and stress conditions (Halliwell et al. 1995; Begcy et al. 2012) directly impacting complex aspects of seed quality including nutritional components, seed size, water potential and dry matter content (Bailly 2004, 2019; Corbineau 2012). In seeds, antioxidants are synthesized throughout the developmental program (i.e., histodifferentiation, reserve accumulation, and late maturation) and are thought to protect against stresses by quenching reactive oxygen species (ROS) and free radicals (Bailly 2004; Chen et al. 2016; Adetunji et al. 2021). For instance, antioxidants balance ROS and free radicals generated in hydration-associated active metabolism from the initiation of embryogenesis through germination by acting as free radical scavengers (Pehlivan 2017). Seed-based antioxidants also influence the hydration process through the action of seed coat phenolics, which may offer a barrier to rapid imbibition (Ross et al. 2010). They also regulate cellular redox balance preventing oxidative damage during the imbibition process (Bailly 2004). Antioxidants also substantially buffer the damage from ROS and other radicles generated by desiccation stress and lipid oxidation during seed storage (Groot et al. 2015; Carta et al. 2018; Adetunji et al. 2021; Stegner et al. 2022), Furthermore, antioxidants may be linked to maintaining seed persistence and viability in the soil seed bank (Long et al. 2015).

At an ecological level, plants adapt to prevailing environmental conditions such as precipitation, relative humidity, solar radiation, and temperatures, allowing normal plant development and reproductive success (Saatkamp et al. 2019). Such adaptions result in the development of fitness-related traits vital to surviving harsh conditions such as drought and cold stress (Capblancq et al. 2022; Eckert and Neale 2022). Studies have indicated a strong correlation between environmental conditions and plant traits. For example, the influence of geographical location on seed germination has been observed in sea oats (Pérez and Kane 2017). At the same time, environmental conditions during plant growth, flowering, and seed development has been found to influence subsequent seed germination and seedling response to oxidative stress (Nguyen et al. 2021).

Seeds traits including seed dormancy and vigor as well as resilience to environmental stresses have been suggested to vary broadly across a latitudinal gradient (Baskin and Baskin 2014; Pérez and Kane 2017; Moreira et al. 2020; Zhou et al. 2021). Using sea oat populations distributed across the US Atlantic and Gulf Coastlines, we investigated the functional dynamics of antioxidant pools in sea oats seeds collected from a continental-scale spatial distribution. We categorized the sea oat populations into four groups based on their phylogeographic classification (Franks et al. 2004). Then, we measured their antioxidant levels hypothesizing possible interactive links of the environmental conditions to the antioxidants levels and seed quality. We compared antioxidant levels from each population to 10-year climate data (precipitation, and temperature) and various ecological classification systems of the study sites. We further tested the potential role of the antioxidant pool on seed quality by correlating antioxidant levels to germination ability.

## MATERIALS AND METHODS

### Plant material collection and seed lot processing

We collected one to three sea oats (*Uniola paniculata*) panicles from no less than 30 widely spaced plants across 18 sites along the US Atlantic and Gulf of Mexico (referred to hereafter as Gulf) coasts in October 2019 (Fig. 1A, Table 1). We spread panicles in a single layer on tarps within a non-climate-controlled warehouse and circulated air over the panicles with an electric fan. We allowed panicles to dry for three days, then hand-stripped spikelets and threshed caryopses (referred to hereafter as seeds) from spikelets with a laboratory debearder (Heavy Duty Batch Debearder, DB3001-600, Mater Seed Equipment, Corvallis, OR). We conditioned seeds further by passing all lots through an air-screen cleaner (Clipper Office Tester, A.T. Ferrell Co. Inc., Corvallis, OR) then an air-column separator (Oregon Seed Blower, Hoffman Manufacturing Inc., Corvallis, OR). Initial germination for fresh seeds ranged from 91 to 100 % on a viable seed basis. We stored remaining seeds in the lab for two and half years (∼ 23 °C, 30-50% RH).

**Figure 1.**
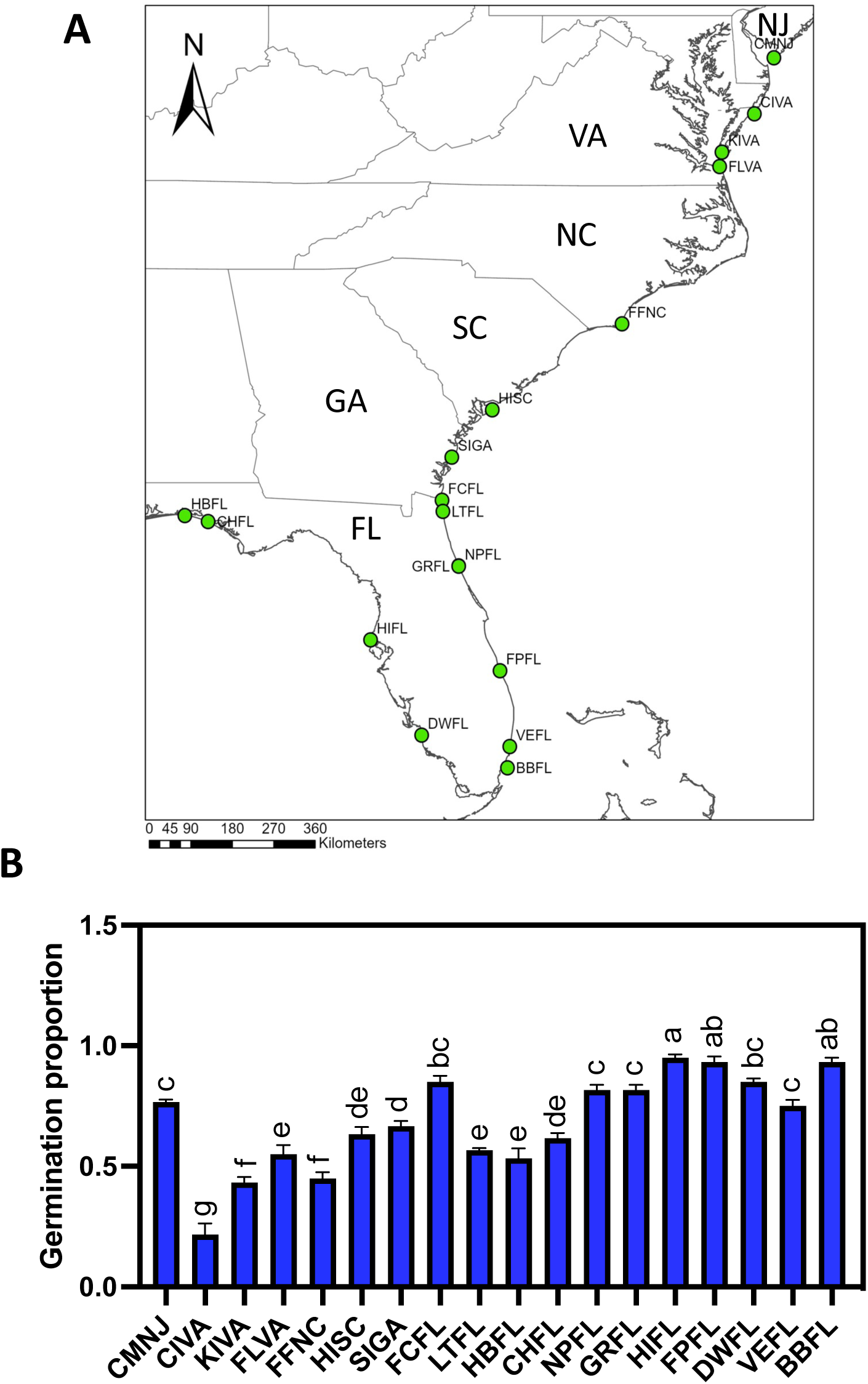
Sea oats seed collection sites and germination from 18 populations collected in the US Atlantic and Gulf Coastlines. (A) Continental-scale map showing the collection sites. North peninsula State Park and Gamble Rogers State Park are close together and appear as a single location. (B) Sea oat seed germination. Seed germination was based on the portion of viable seeds, germination at 28 days, and positive 1% Tetrazolium (TZ).

**Table 1.**
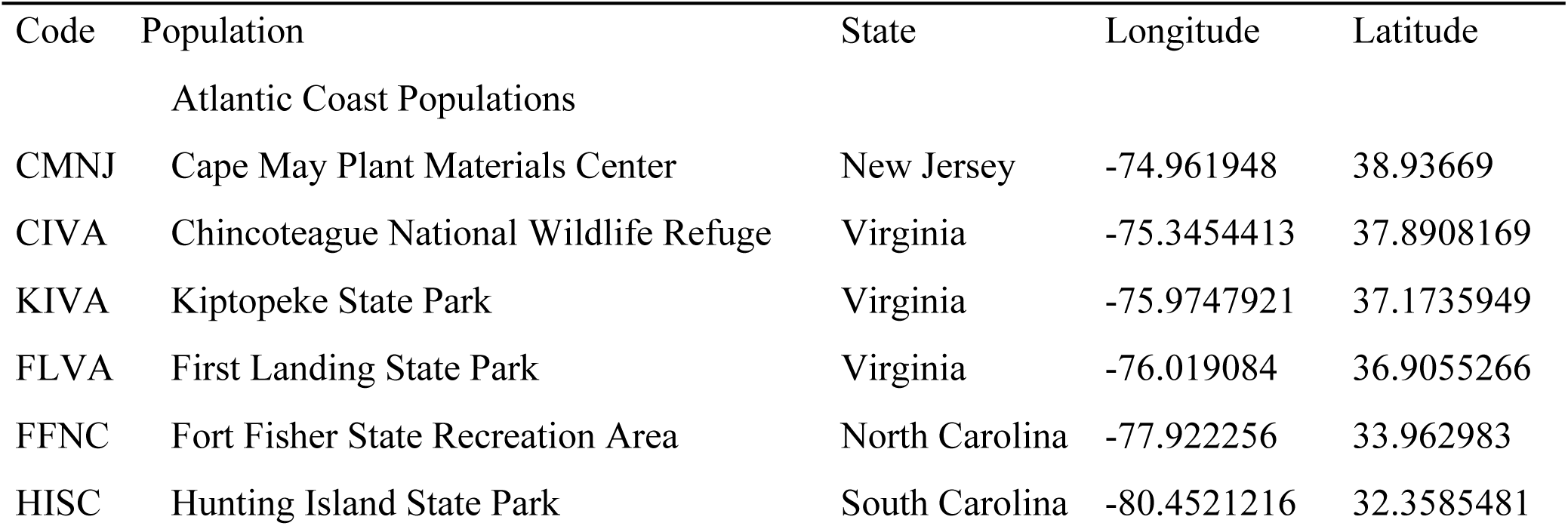
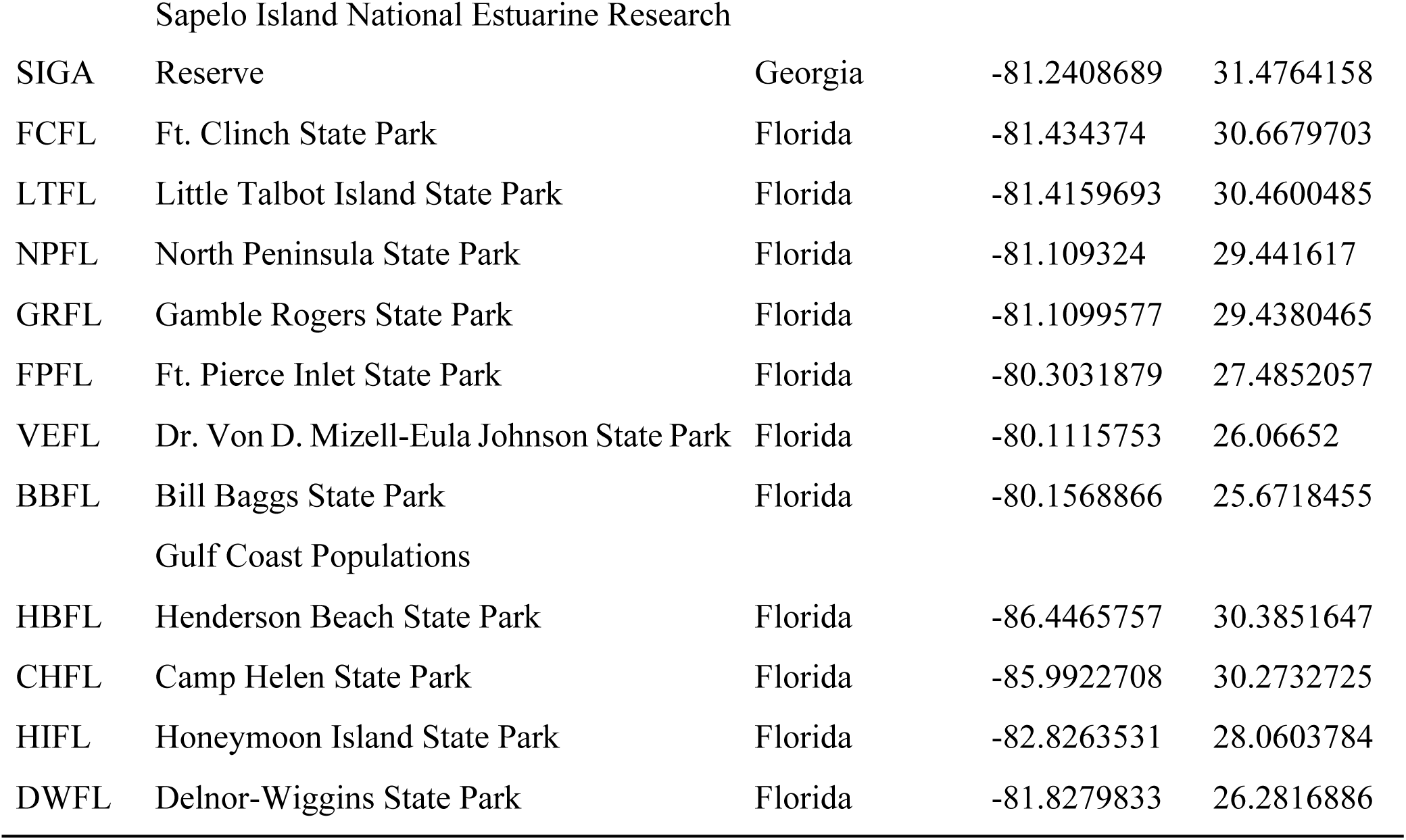
Sea oats seed collections sites from the USA Atlantic and Gulf Coastlines.

### Seed imbibition and germination assays

We randomly selected 200 seeds from each population for imbibition and germination assays. We surface sterilized tested seeds by placing them in 250 mL glass conical flasks with sufficient ethanol (50% v:v) to cover all the seeds (ca., 40 mL). The flasks were transferred to a benchtop shaker and agitated at 200 rpm for 10 minutes. This was followed by a double rinse (i.e., 3 min each) with 100 mL of sterilized distilled H_2_O followed by adding 40 mL of 25% sodium hypochlorite and incubation on a benchtop shaker at 200 rpm for 5 minutes, then another double rinse with 100 mL of sterilized, distilled H_2_O (i.e., 3 min each) before using them in imbibition and germination assays (Begcy et al. 2018).

We performed seed imbibition experiments by placing fifteen seeds on blotter paper (Steel Blue, Anchor Paper Co., St. Paul, MN) moistened with 30-50 mL distilled H_2_O containing 0.2% solution of a broad-spectrum biocide (Plant Preservative Mixture, Plant Cell Technologies, Washington, D.C.) contained within polystyrene germination boxes (156C, Hoffman Manufacturing, Corvallis, OR). We then transferred boxes to an incubator (I30-VL, Percival Scientific, Perry IA) set to 35/25 °C (day/night) and a 12-h photoperiod. Illumination coincided with the day temperature. We allowed seeds to imbibe under these conditions for 36 h, after which we froze the seeds in liquid nitrogen and stored them at −80 °C for use in the determination of Trolox Equivalent Antioxidant Capacity (TEAC) as described below.

We used three to four samples of 25 seeds in our germination experiments that were conducted under the same conditions described for the imbibition experiments. However, germination assays continued for 28 days. We inspected the germination boxes daily and counted germination when seeds exhibited 2 mm of radicle and coleoptile emergence (Begcy et al. 2018). We removed germinated seeds and any seeds showing signs of fungal contamination. We assessed the viability of any remaining, non-germinated seeds after day 28 using a tetrazolium stain (Peters, J. and Lanham, B. 2000). We analyzed embryo staining patterns at 2.5 to 6.4× and report germination on a viable seed basis (Supplementary Fig. S1A-B).

### Seed water relations

We measured post-storage seed water potential, using three technical replicates, with a dewpoint potentiometer (WP4C, Decagon Devices, Pullman, WA). The potentiometer was calibrated before each measurement day according to manufacturer recommendations. We measured seed water potential at 20 °C in precise mode. In this procedure, we used about 200 seeds to cover the bottom of the cuvette completely. Seed moisture content was calculated by recording the fresh mass on 20 seeds in five replicates for each of the 18 populations studied. We then dried these seeds in a forced-air oven (Blue M, Illinois, USA) at 103 °C for 17 hours and report water content on a dry mass basis (g H_2_O · g dry seed mass^-1^ [g g ^-1^]).

### Climate and ecological data

We obtained climate data for the ten years before our seed sampling (2009-2019) using the PRISM application (Hart and Bell 2015). The climate data used in this study included minimum (Tmin), mean (Tmean) and maximum (Tmax) temperatures, mean dew point temperature (Tdmean), total precipitation (ppt), and daily maximum vapor pressure deficit (Vpdmax). All data obtained were filtered at 4 km resolution using the location coordinates for sites corresponding to each of the 18 sea oat populations (Fig. 1A, Supplementary Table S1). Data were extracted using the prism R package (Hart and Bell 2015).

First, we explored seasonal precipitation and temperature patterns along the latitudinal gradient where the sea oats populations were collected. We organized our analysis into groups of three months, roughly representing the four seasons (Winter, Spring, Summer, and Autumn).

Ecological descriptors comprised the: U.S Environmental Protection Agency (US EPA) ecoregion zoning (EPA3 and EPA4) (Supplementary Table S2), U.S. Department of Agriculture-Natural Resource Conservation Service (USDA-NRCs) and Food and Agriculture Organization of the United Nations (FAO) soil classification (Supplementary Table S3), American Horticulture Society (AHS) heat zoning (Supplementary Table S4), USDA plant hardiness zoning (Supplementary Table S5). We also used sea oat phylo-geography data as population grouping factors (Franks et al. 2004 Supplementary Table S6).

### Antioxidant analysis

We estimated the levels of Trolox Equivalent Antioxidant Capacity (TEAC) in seeds using the modified ABTS/TEAC assay (Re et al. 1999). ABTS/TEAC determines the TEAC per gram of sample. ABTS (2,2’-Azino-bis (3-ethylbenzothiazoline-6-sulfonic acid) reagent was prepared by dissolving 2.74 mg of ABTS in 1 mL of sterile, distilled H_2_O for each reaction, including the standards. Then, 300 mg of MnO_2_ was added and mixed using a magnetic stirrer for 20 minutes. The mixture was filter sterilized through a 0.45 μm syringe filter. For calculations, a spectrophotometer (Scientific Genesys 10S UV-VIS Spectrophotometer, Thermo Scientific, Madison, USA) was zeroed at 734 nm with 100 μL PBS added to 1 mL of sterile distilled H_2_O. Absorbance readings were adjusted with 5 mM PBS to obtain a value of 0.700. We added 6.26 mg of Trolox (6-hydroxy-2,5,7,8-tetramethylchroman-2-carboxylic acid) to 50 mL of 5mM PBS then mixed the reagents by sonication for 20 minutes to obtain 0.5 mM Trolox standard solution. Then, 1 mL of aliquots were prepared in cryovials and stored in a freezer at −20 °C. During the assay, a 1 mL cryovial containing 0.5 mM Trolox was obtained from the freezer and allowed to thaw. We re-dissolved the solution by sonication for 10 minutes. Then, dilutions for five triplicate standards were obtained at 0, 11, 22, 33, and 44 μl of 5mM Trolox corresponding to TEAC concentrations of 0, 5, 10, 15, and 20 μmol/L, respectively.

Antioxidants for TEAC estimation were extracted from seeds using a modified hydrophilic method (Re et al. 1999). Samples were finely ground using liquid nitrogen. Approximately 10 mg/sample was placed into 10×75 mm reaction vials. Triplicates were used per sample per reaction. TEAC was extracted by adding 5 mL of 75% aqueous methanol, followed by stirring under a nitrogen gas stream of 2.0 Pascals at 30 °C for 60 minutes. Then, the obtained product was centrifugated at 2000 × g for 5 minutes to get a supernatant containing the extracts. A second extraction step was conducted by adding 2 mL of 75% aqueous methanol and covering the reaction vials. Then, each vial was vortexed and centrifugated at 2000 × g for 15 minutes. Supernatants of the first and second reactions were combined. Concurrently, 10-20 mg of finely ground samples were prepared in oven-drying foil pans and dried at 103 °C for 17 hours to obtain dry mass.

Extracts were re-suspended into 10 mL of 75% methanol. Each 100 μL of sample extract was mixed with 1 mL of 5 mM ABTS solution, vortexed for two mins, and placed in the spectrophotometer cuvette before reading the absorbance at 734 nm. Three biological replicates and three technical replicates were used.

### Data analysis

We subjected data on seed water potential, moisture content, TEAC, and germination percentage to an analysis of variance (ANOVA) to estimate differences among populations. We utilized linear and polynomial regression to evaluate seed germination and TEAC levels against geographical, climatic, and ecological patterns. We also calculated Cohen’s f^2^ to standardize parameter effect size and used the following guidelines for interpretation: f^2^ ≥ 0.02 represents a small effect; f^2^ ≥ 0.15 medium effect; and f^2^ ≥ 0.35 large effect (Cohen 1988). Statistical analyses were performed using the R software/environment 1.3. Differences were considered statistically significant at p-value < 0.05 (Kim et al. 2021).

## RESULTS

### Impact of the latitudinal gradient on seed physiology in sea oat populations along the US Atlantic and Gulf Coastlines

To test the hypothesis of seed related traits variation across a wide latitudinal gradient (Baskin and Baskin 2014; Pérez and Kane 2017; Moreira *et al*. 2020; Zhou et al. 2021), we collected and germinated eighteen sea oat populations distributed across the US Atlantic and Gulf Coastlines (Fig. 1A). First, we categorized the sea oat populations into four groups based on their phylogeographic classification (Franks et al. 2004). Seed germination ranged from 20% (CIVA) to 95% (HIFL) (Fig. 1B). Seeds from 39% of the populations registered germination >80%, most of the populations displaying this level of germination were from Florida (Fig. 1B). However, there were instances of poor germination (i.e., < 50%) for other populations, especially from the northern latitudes of the Atlantic coastline (Fig. 1B). Interestingly, seeds collected from populations in the southern US displayed germination percentages > 55% (Fig. 1B).

To further understand differences in seed physiology that may influence variation of germination, we quantified seed moisture content and water potential (Fig. 2A-B). Seed moisture content was relatively similar across the latitudinal gradient and ranged from 0.1064 to 0.1129 g g^-1^. Seeds with lower moisture content (< 0.1100 g g^-1^) comprised KIVA, FFNC, SIGA, GRFL, NPFL, and HBFL (Fig. 2A). While seeds collected from LTFL and CIVA displayed the highest water contents observed of 0.1129 and 0.1130 g g^-1^, respectively. The seeds with the lowest moisture content were collected from Florida. They included GRFL and NPFL at 0.1066 and 0.1064 g g^-1^, respectively (Fig. 2A). Sea oats seed water potential was also variable and ranged from −89.01 to −100.84 MPa (Fig. 2B). Patterns of water potential did not necessarily track with patterns of water content as some populations displaying lower water content expressed higher water potential values. Conversely, other populations displaying higher water contents displayed lower water potentials (Fig. 1B and 2A-B). However, differences in seed water relations could not explain germination variation across populations.

**Figure 2.**
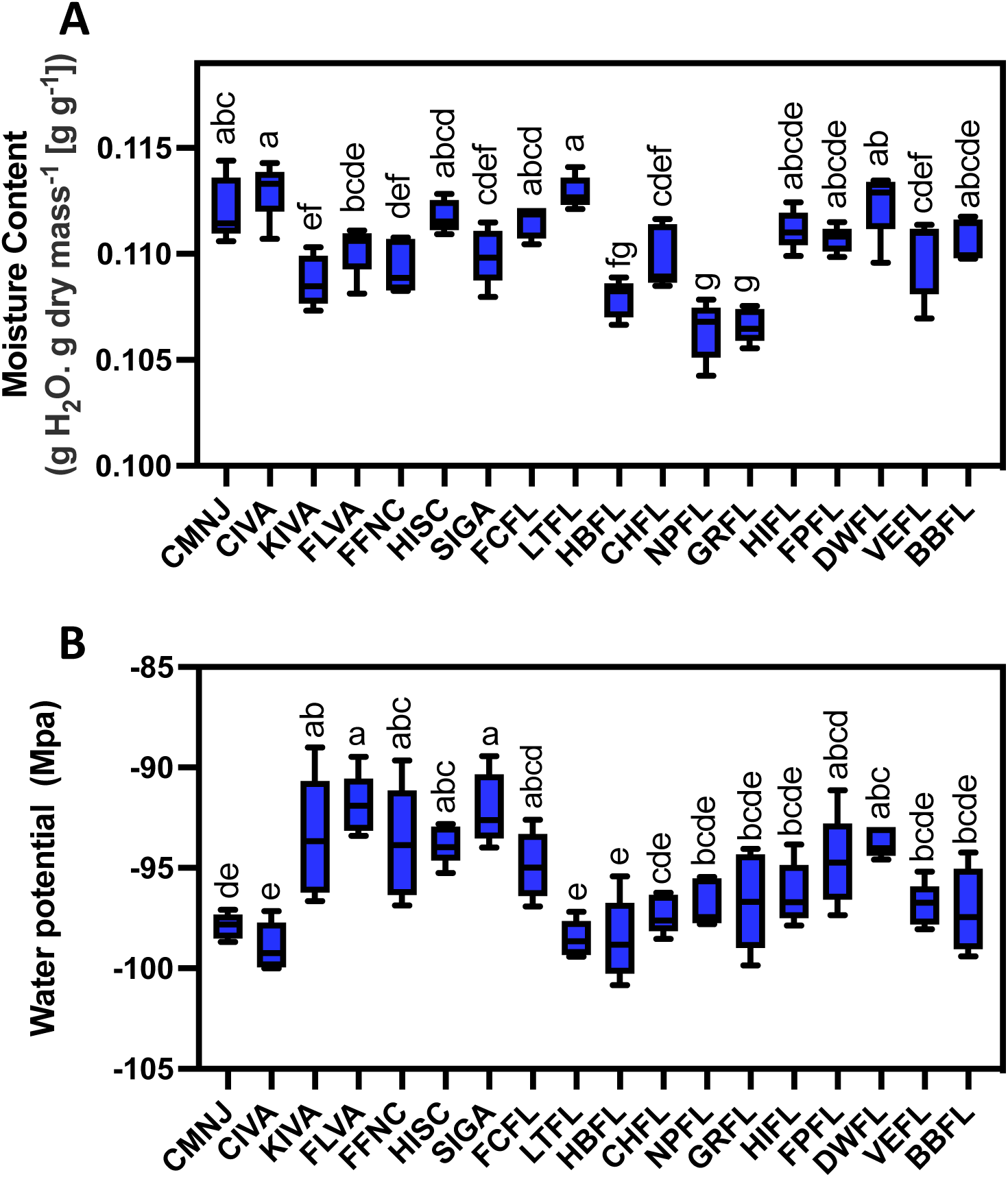
Limited variation on moisture content and water potential of sea oats seeds collected across the US Atlantic and Gulf Coastlines. (A) Moisture content (g H2O g-1) and (B) water potential (Mpa).

To characterize the extent to which location and environmental conditions influenced germination of sea oats seeds, we examined how germination related with latitude and temperature zoning (Fig. 3; Supplementary Tables S4 and 5). We observed a latitudinal pattern of germination response, with regions closer to the equator having higher germination while populations located further from the equator displayed reduced germination capacity (R^2^ = 0.50), indicating a significant latitudinal influence on germination ability (Fig. 3A). In addition, germination was lower in zones with fewer heat days above 30 °C but increased as the number of heat days increased (Fig. 3B). Similarly, germination was lowest in zones with lower average minimum winter temperatures but increased as this average rose (Fig. 3C). We also calculated Cohen’s f^2^ to standardize parameter effect sizes and found a large effect of latitude and temperature zoning on germination capacity (Fig. 3).

**Figure 3.**
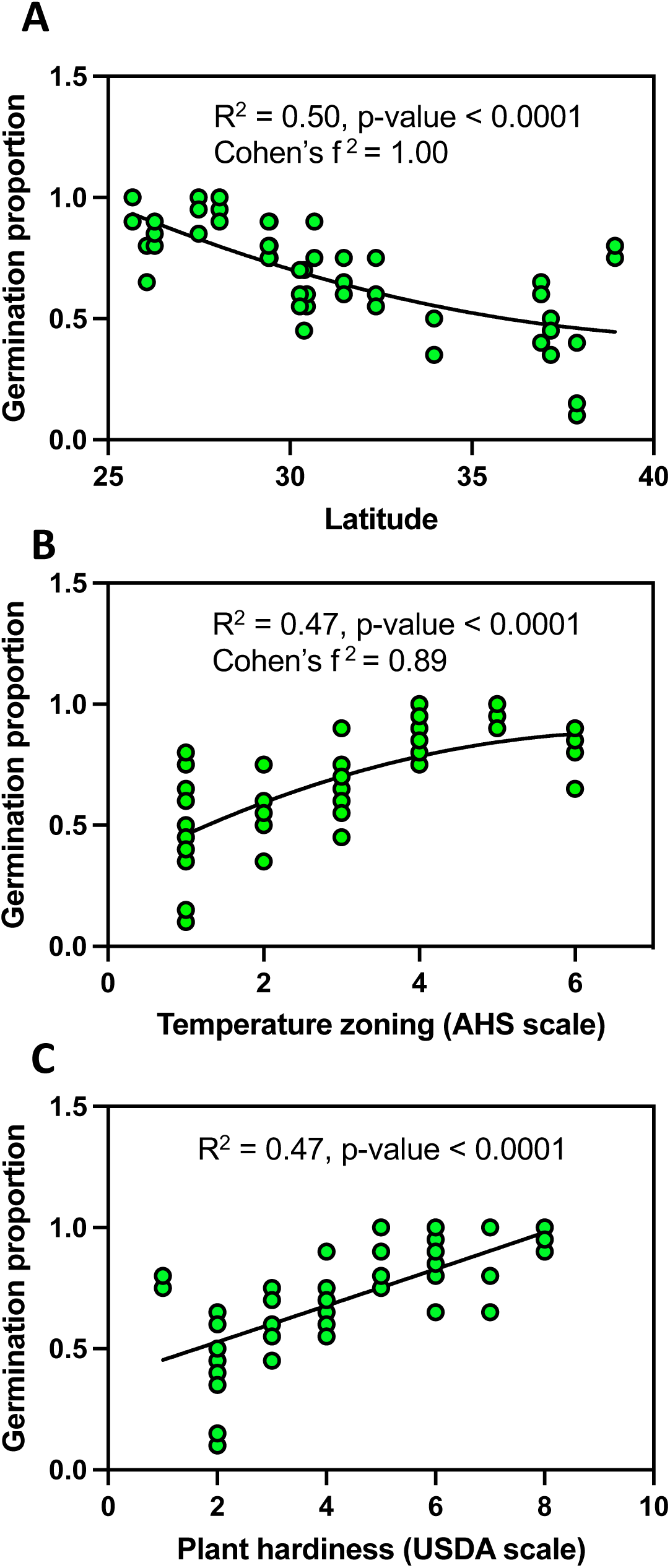
Influence of location and environmental zoning by temperature factors on the germination of sea oat seeds. (A) latitude, (B) AHS temperature zoning, and (C) USDA plant hardiness zoning. The AHS temperature zoning is based on days at temperatures above 86oC, covering 45-60 days (1), 60-90 days (2), 90-120 days (3), 120-150 days (4), 150-180 days (5) and 180-210 days (6) per year. USDA plant hardiness scale is based on vegetation exposure to cold conditions in winter, at levels of 1(0-10F), 2(10-15F), 3(15-20F), 4(20-25F), 5(25-30F), 6(30-35F), 7(35-40F) and 8(40-45F). The germination pattern model follows Y = B0 + B1 + B2×2.

### Weather characteristics of sea oat seeds collection sites along the US Atlantic and Gulf Coastlines

To explain which factors could have contributed to the observed differences in seed traits across all sea oats populations (Fig. 1-3), we first looked at precipitation and seasonal mean temperature patterns for sea oats collection sites along the US Atlantic and Gulf coastlines in a 10-year period prior to seed sampling (2009-2019). Precipitation patterns across the latitudinal gradient were similar in winter and spring, averaging 50 to 150 mm (Fig. 4A). However, in summer, southerly latitudes displayed higher precipitation (>150 mm) than northerly latitudes (<150 mm). In Autumn, the southerly latitudes maintained higher precipitation in September and October (>100 mm) before dropping below 100 mm in November. Alternatively, precipitation increased considerably in northern latitudes to > ca.100 mm in September and October but remained lower (<100 mm) in November (Fig. 4A).

**Figure 4.**
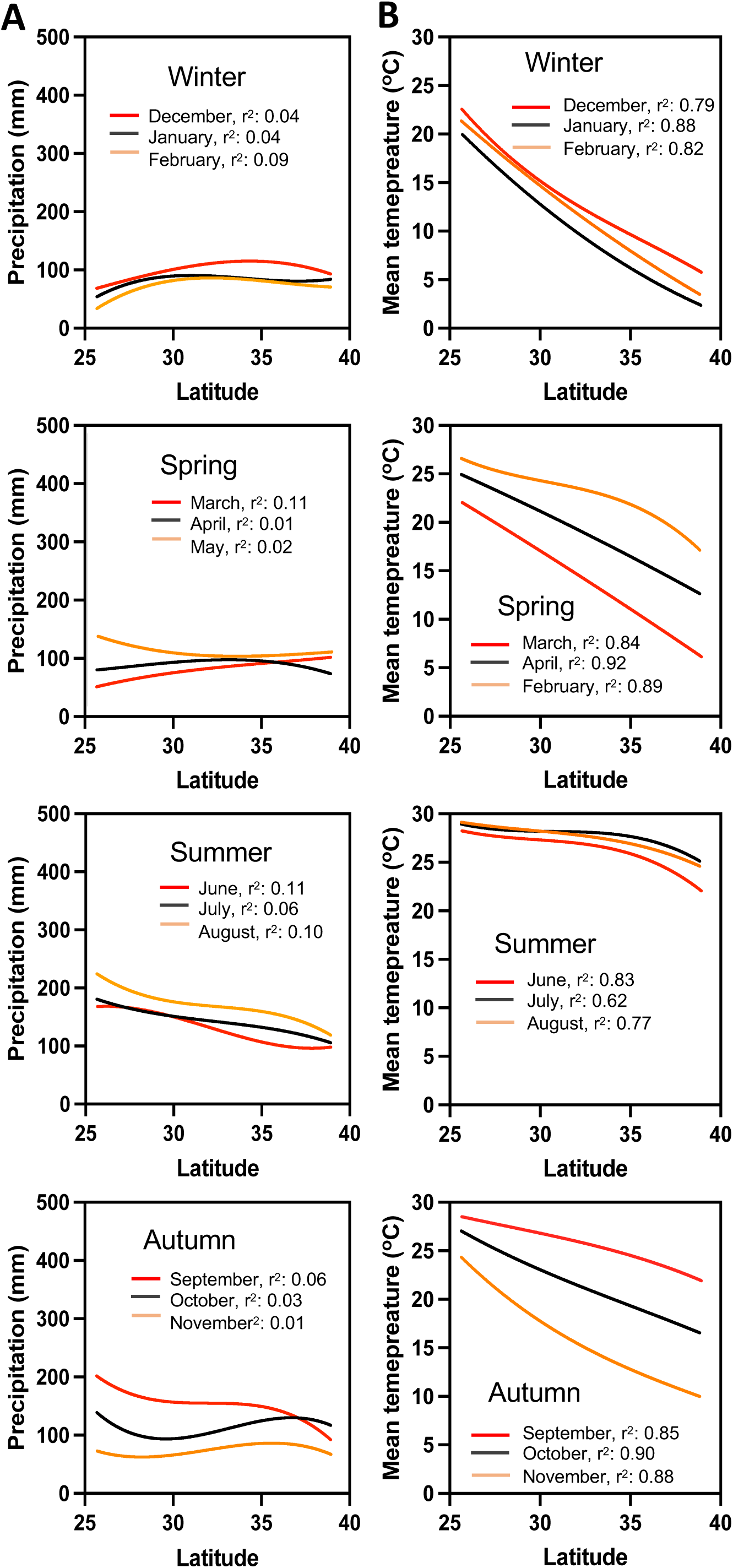
Environmental data patterns for the sites of sea oats collection from the US Atlantic and Gulf coastlines in a 10-year period prior to seed sampling (2009-2019). (A) Precipitation and (B) seasonal mean temperature. Model for precipitation and temperature patterns follow the function Y = B_0_ + B_1_ + B_2_x^2^ + B_3_x^3^.

Temperature patterns were similar across all seasons (Fig. 4B). However, higher temperatures were evident in southern latitudes compared to the northern ones during the Winter, Spring, and Autumn (Fig. 4B). Summer temperatures varied less across all the latitudes in which the sea oats populations were analyzed (Fig. 4B). Mean temperatures across all latitudes remained above 22 °C throughout June, July, and August (Fig. 4B). Both the maximum vapor pressure deficit and the mean dew point had a similar pattern, with higher values in the southern latitudes and lower values in the northern latitudes in most seasons (Winter, Spring, and Autumn), except in summer season in which there were no differences between southern and north latitudes (Supplementary Fig. S2-3).

### Latitude-dependent environmental conditions influence TEAC concentration in sea oats seeds

Since our climate data pointed to precipitation and temperature as main drivers of germination differences across a latitudinal gradient, we used antioxidant levels in sea oats seeds as a proxy to explore a possible link of latitude-dependent environmental conditions with antioxidant accumulation and germination. First, we defined baseline TEAC values obtained from mature ungerminated seeds before imbibition across all sea oats populations. We found that TEAC levels from our collected seeds ranged from 0 to 12 μmol/g DW (Fig. 5A). For instance, while seeds collected in CMNJ, CIVA, KIVA, VEFL, BBFL, HIFL DWFL and CHFL showed TEAC levels higher than 4 μmol/g, the remining populations showed lower values than that (Fig. 5A). Interestingly, in general, seeds from sea oat populations collected either from the most northern or southern locations showed higher antioxidant capacity than the ones collected in intermediate latitudes (Fig. 5A). These results suggest an influence by extreme conditions experienced at northern or southern US latitudes. To further explore whether antioxidant capacity was maintained during the initiation of germination, we imbibed seeds for 36 h and quantified their TEAC levels (Fig. 5B). Contrary to the earlier patterns of baseline antioxidant levels, there was wider population variability across the geographical distribution of sea oats (Fig. 5A-B). The change in TEAC levels between dry and imbibed seeds showed considerable variation (Fig. 5C). For instance, seeds from some latitudes displayed a drastic drop in TEAC levels, especially CMNJ and KIVA (Fig. 5C), and those with significantly minimal changes comprised KIVA, FPFL, BBFL, DWFL, HBFL, and CHFL (Fig. 5C). However, some seeds registered a significantly positive change in TEAC > 5 μmole/g DW comprised of FLVA, FFNC, HISC, SIGA, FCFL, LTFL, GRFL, NPFL and VEFL (Fig. 5C).

**Figure 5.**
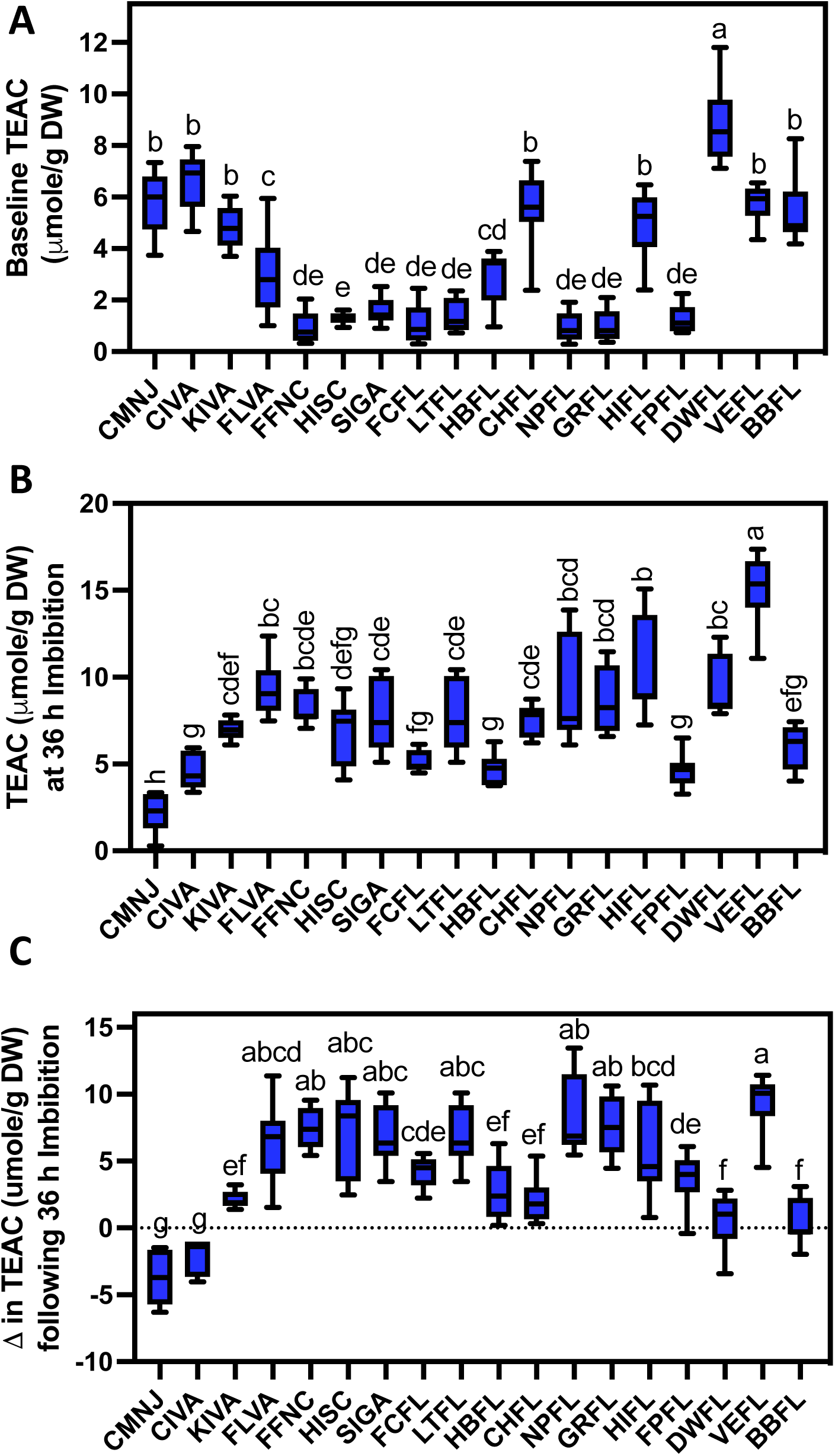
Baseline antioxidant levels of dry seeds and after 36 h of imbibition collected from a continental-scale spatial distribution. Baseline TEAC on (A) dry seeds, (B) after 36 h of imbibition, and the (C) difference in TEAC between dry seeds and after 36 h of imbibition.

### Geographic and environmental patterns in relation to seed traits in sea oats from US Atlantic and Gulf coastlines

To assess the impact of latitude on antioxidant capacity, we used baseline seed TEAC concentrations and compared these with the latitudinal range where seeds were collected. In general, antioxidant levels in dry seeds displayed a concave parabolic pattern when evaluated against latitude, heat, and plant hardiness zones (Fig. 6A-C). Then, we compared antioxidant capacity levels obtained 36 h after imbibition with the latitudinal zones (Fig. 6D). We observed higher TEAC levels near the equator and lower levels in northern latitudes. Surprisingly, relationships between TEAC and AHS heat or USDA plant hardiness classifications showed similar patterns when TEAC data obtained from seeds imbibed after 36h (Fig. 6E-F). Zones with a greater number of hot days (Fig. 6F) and less freezing conditions (Fig. 6E) displayed higher TEAC values. We found large effect on TEAC obtained from dry seeds compared against latitude (Fig. 6A) and temperature zoning (AHS heat) (Fig. 6B) as well as on temperature zoning in 36 h imbibed seeds (Fig. 6E). We found medium effect size when TEAC obtained from dry and imbibed seeds were compared against USDA plant hardiness (Fig. 6C, F) and imbibed seeds against latitude (Fig. 6C)

**Figure 6.**
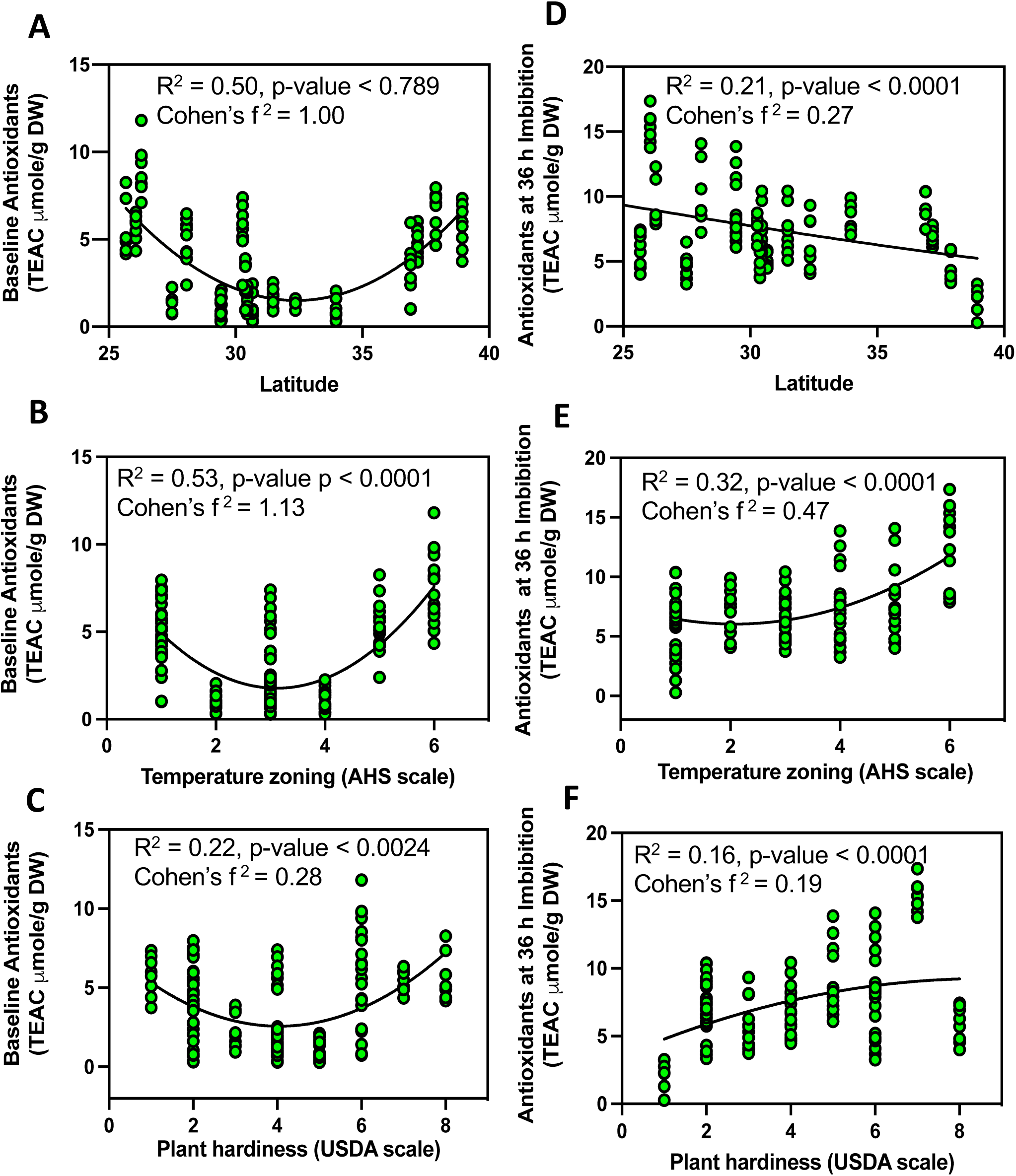
Influence of location and environmental zoning by temperature factors on antioxidant levels. Relationship of TEAC in dry seeds (A-C) and after 36 h of imbibition (D-F). (A-D) Latitude, (B-E) AHS temperature zoning, and (C-F) USDA plant hardiness zoning. The AHS temperature zoning is based on days at temperatures above 86°C, covering 45-60 days (1), 60-90 days (2), 90-120 days (3), 120-150 days (4), 150-180 days (5) and 180-210 days (6) per year, while the USDA plant hardiness scale is based on vegetation exposure to cold conditions in winter, at levels of 1(0-10F), 2(10-15F), 3(15-20F), 4(20-25F), 5(25-30F), 6(30-35F), 7(35-40F) and 8(40-45F). The TEAC patterns model follows the function Y = B0 + B1 + B2^x2^.

### Role of the environment and seed traits on the baseline germination and antioxidants in sea oats from the US Atlantic and Gulf coastlines

No clear relationships between germination and baseline TEAC or TEAC after 36 h of imbibition were evident when considering the geographical range of sea oats used in this study (Supplementary Fig. S4A-B). Therefore, we analyzed relationships between germination and TEAC using subsets of data representing the extreme temperature regions where high TEAC levels had been observed. Similarly, no clear relationships existed between germination and TEAC after 36 h of imbibition for seeds originating from colder regions (>33N) (Fig. 7A-B). However, seeds from warmer regions (< 30N), where high seed quality was expected, exhibited a significant negative linear relationship of TEAC levels in imbibed seeds after 36 h showing large effect sizes according to Cohen’s f^2^ coefficient (0.39, 0.43, Fig. 7C-D, respectively).

**Figure 7.**
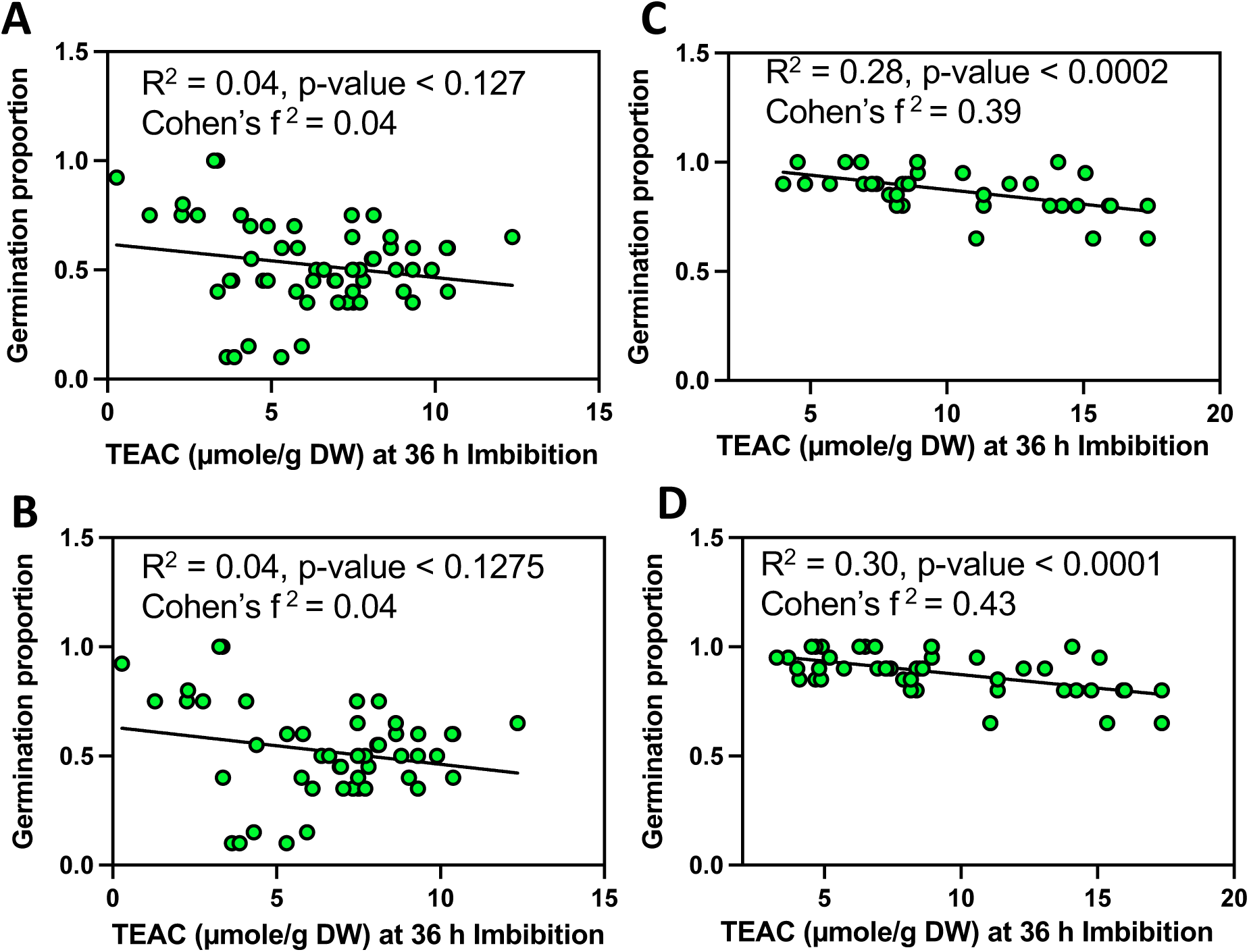
Influence of TEAC levels on germination of sea oats seed populations. (A) AHS Temperature zone ≤ 2; (B) AHS Temperature zone > 4; (C) USDA plant hardiness zone ≤ 3 and (D) zone ≥ 6. These zones correspond to latitudinal locations of >33N (A and C) and < 30N (B and D). The AHS temperature zoning is based on days at temperatures above 86oC, covering 45-60 days (1), 60-90 days (2), 90-120 days (3), 120-150 days (4), 150-180 days (5) and 180-210 days (6) per year, while the USDA plant hardiness scale is based on vegetation exposure to cold conditions in winter, at levels of 1(0-10F), 2(10-15F), 3(15-20F), 4(20-25F), 5(25-30F), 6(30-35F), 7(35-40F) and 8(40-45F).

Finally, we compared the influence of latitude and temperature patterns on TEAC levels on dry seeds and seeds imbibed for 36 h (Fig. 8). We observed that the minimum and maximum temperatures during the seed development period exhibited high TEAC levels in seeds collected from northern or south latitudes (Fig. 8A). As such, the minimum and maximum temperatures exhibited latitudinal patterns of distribution (R^2^ of 0.22 and 0.20, respectively). Contrary, 36 h of imbibition resulted in lower TEAC levels on seeds collected on northern sites (Fig. 8B).

**Figure 8.**
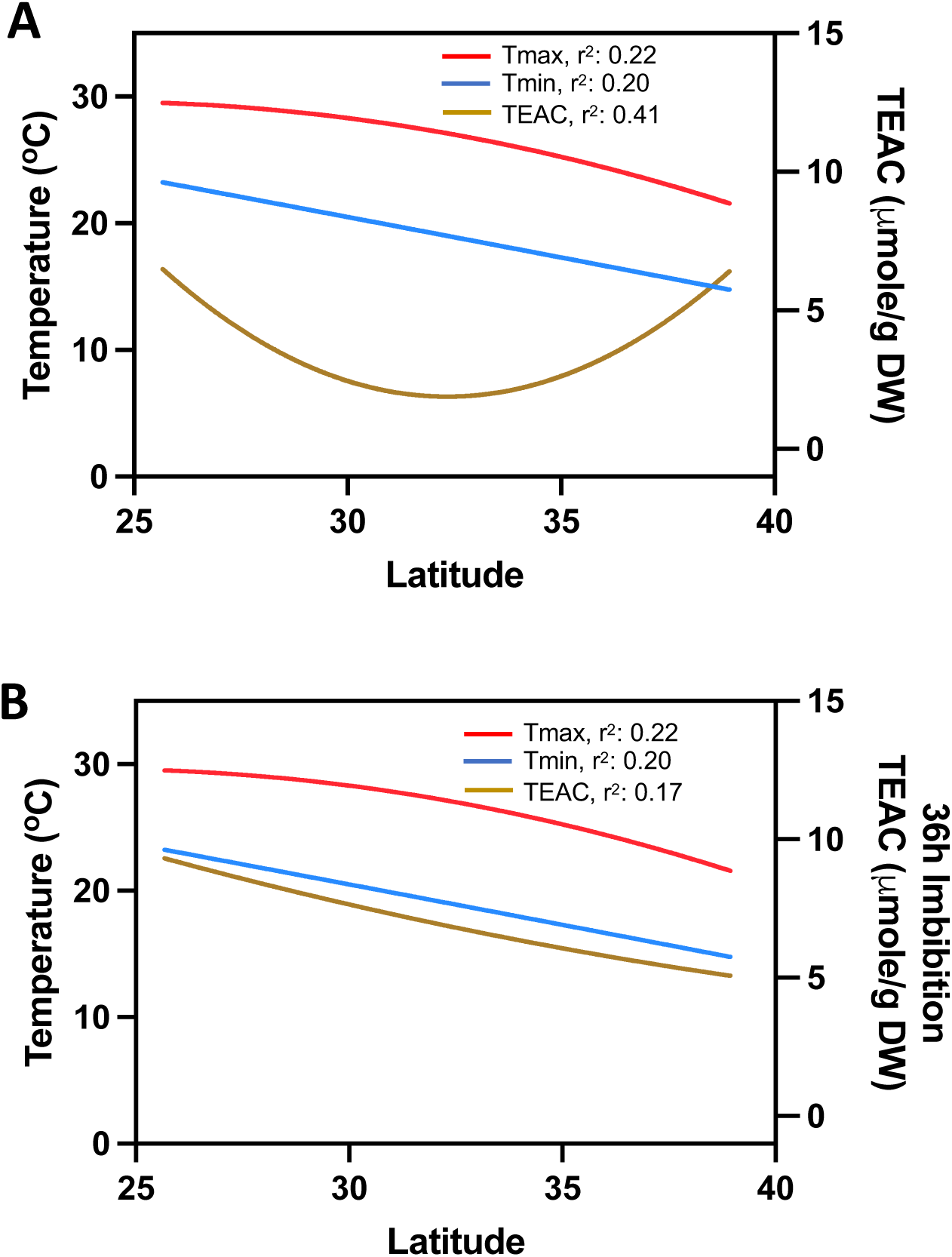
Influence of location and latitudinal temperature patterns on baseline antioxidants in sea oats. Tmax: maximum temperatures recorded during the day. Tmin: minimum temperatures recorded during the day in relation to (A) baseline TEAC and (B) after 36 h of imbibition. Temperature and TEAC patterns models were calculated using the equation Y = B_0_ + B_1_ + B_2_x^2^.

## DISCUSSION

Sea oats are the primary native dune grass used to restore and stabilize beaches and dunes across the southeastern United States. Commercial nurseries have propagated sea oats using field-collected seeds to retain the genetic integrity of source populations. We investigated the effect of climate conditions on sea oats harvested across a wide latitudinal range that spanned more than 13 ° of latitude (Table 1). Our results show that sea oats seed germination was higher from the sites closer to the equator but decreased as the distance from the equator increased (Fig. 1B). The observations of comparative germination with respect to distance from the equator indicated that the sea oat seeds are of high quality in terms of germination percentage from warmer tropical regions compared to seeds from colder areas. Similar patterns of seed germination were observed between populations located in Southern Italy in comparison with a Spanish core population of *Lavandula multifida L.,* a plant species present in the Western Mediterranean Basin (Panuccio et al. 2018).

### Environmental and climate influences on seed quality, antioxidants, and implications on seed germination

Antioxidant activity plays a key role in various events of seed life. Antioxidants are produced throughout the seed developmental program and during the germination process. These molecules occur within metabolically active cells, but also in dry tissues, with physiological activity depending on seed moisture status (Vertucci and Farrant 1995; Walters *et al*. 2002; Ballesteros *et al*. 2020). Water potential measurements of dry seeds tested in this study suggest that seeds from all populations fall into the same hydration level (i.e., Hydration level II, ca. −15 to 190 MPa (Vertucci and Farrant 1995; Walters et al. 2002, 2005) which was like previous studies (Pérez and Kane 2017). Hydration level II represents a cellular physiological state characterized by enzyme-mediated catabolic reactions, oxidative and peroxidative processes, and formation of free radicals and reactive oxygen species. Existing pools of molecular antioxidants synthesized during the seed developmental program are thought to play a protective role by quenching reactive oxygen species and free radicals in dry seeds (Vertucci and Farrant 1995; Walters et al. 2002, 2005). We did not observe a clear relationship between TEAC concentration and baseline seed water potential in this study. Nonetheless, seeds from all populations display antioxidant capacity that may be sufficient to mediate aging associated deteriorative reactions to some degree during storage under non-optimal conditions of temperature and relative humidity (Walters et al. 2005). Moreover, it is well documented that environmental conditions directly impact seed quality as well as their desiccation tolerance, nutrient level, storage potential and the accumulation of antioxidants and other related secondary metabolites (Bewley et al. 2013; Pehlivan 2017; Ellis 2019; Malovichko et al. 2021). We observed differences in the accumulation of antioxidant levels in dry seeds compared to imbibed seeds across all populations (Fig. 5-6). Increased antioxidant activity during seed germination is vital in signaling and ROS homeostasis (Bailly 2004; El-Maarouf-Bouteau and Bailly 2008). High germination capacity associated with moderate changes in TEAC levels after initial hours of imbibition is expected to be a characteristic of good-quality seeds because it is linked to low lipid peroxidation (Rogozhin et al. 2001). Interestingly, in our study, seeds from warmer regions exhibited limited fluctuations in TEAC levels after imbibition. Therefore, sharp increases in TEAC levels for seeds collected from populations located in warmer regions might be an adaptative response to climate conditions that directly impact their germination potential. These results agree with the elevated antioxidant levels (α-tocopherol and tocotrienols) found in flax (*Linum usitatissimum*) (Oomah et al. 1997) and in barley (*Hordeum vulgare*) seeds after elevated temperatures (Roach et al. 2018). Similarly, a negative correlation between antioxidants (α-tocopherol and β-tocopherols) was associated with poor longevity for rice seeds from temperate regions (Lee et al. 2019). Therefore, the antioxidant variation may have been conditioned by environmental conditions as reported in other plant species.

### Latitudinal gradients correlate with antioxidant accumulation

We aimed to answer whether latitudinal gradients have an impact on seed trait performance. Our results showed variation on germination and antioxidant accumulation along the latitudinal gradient (Fig. 1B and 6). Interestingly, the latitude-associated geographic distribution of antioxidants in leaves of Bilberry (*Vaccinium myrtillus* L.) was linked to adaptions to prevailing conditions and prevailing abiotic stress associated with a given geographic location (Martz et al. 2010). Similarly, secondary metabolites such as anthocyanidin are strongly controlled by the latitude and geographic origin in *V. myrtillus* fruits (Åkerström et al. 2010). A similar case was reported for blueberries (*Vaccinium ashei* cv. ‘Brightwell’) in higher altitudes with higher concentrations of flavonoids, phenols, proanthocyanidins, and anthocyanins, among other compounds (Zeng et al. 2020). Our data shows that latitude influences seed germination. However, this influence might be as the combinatorial result of the climate conditions. Therefore, further evidence needs to be provided to separate those variables.

## CONCLUSION

In summary, our results show a significant influence on seed physiological performance along a latitudinal gradient in sea oats. Combined with the environmental variables, latitude directly dictates antioxidant accumulation in ungerminated seeds and after imbibition. Antioxidants are among the seed storage components critical for persistence, longevity, storage quality, and viability (Long et al. 2015). Therefore, the involvement of the environment in accumulating these secondary metabolites alongside other storage reserves is critical in seed quality. Knowledge of the interaction of associated environmental factors and their interaction is crucial for plants. It can be helpful in guiding the collection of seed materials and seed quality control for storage and use in conservation efforts.

## SUPPLEMENTARY DATA

Supplementary data are available online and consist of the following:

Table S1. Ten-year (2009-2019) climate data from study sites obtained from PRISM.

Table S2. Environmental data (EPA classification)

Table S3. USDA-NRCS and FAO soil classification for the study sites.

Table S4. American Horticultural Society Plant heat zones.

Table S5. USDA plant hardiness scale.

Table S6. Sea oat population classifications.

Figure S1. Tetrazolium staining of sea oats seeds. A. Positive 1% Tz Test and B-Negative 1% Tz test. The 1% Tz test was conducted on ungerminated seeds following 28 days of germination incubation. Seed were suspended in 1% Tz coved and incubated at 35 °C for 24 h before dissection and imaging.

Figure S2. The seasonal vapor pressure deficit patterns for the sites for the collection of 18 populations of Sea Oat seeds from the US Atlantic and Gulf coastlines, from the 2019 collection. (A) Winter; (B) Spring; (C) Summer and (D) Autumn.

Figure S3. Seasonal mean dew point patterns for the sites for the collection of 18 populations of sea oat seeds from the US Atlantic and Gulf coastlines, from the 2019 collection. (A) Winter; (B) Spring; (C) Summer and (D) Autumn.

Figure S4. Influence of TEAC levels on germination of 18 populations of sea oat seeds of the US Atlantic and Gulf coastlines, from the 2019 collection. A. Baseline TEAC B. TEAC following 36 h of imbibition.

## ACKNOWLEDGEMENTS

We thank Dr. William Hammond for his insightful suggestions on the data analysis and Dr. Tia Tyler for the technical assistance during the seed collection.

## LITERATURE CITED

Adetunji AE, Sershen, Varghese B, Pammenter N. 2021. Effects of exogenous application of five antioxidants on vigour, viability, oxidative metabolism and germination enzymes in aged cabbage and lettuce seeds. South African Journal of Botany 137: 85–97.

Åkerström A, Jaakola L, Bång U, Jäderlund A. 2010. Effects of Latitude-Related Factors and Geographical Origin on Anthocyanidin Concentrations in Fruits of *Vaccinium myrtillus* L. (Bilberries). Journal of Agricultural and Food Chemistry 58: 11939–11945.

Bailly C. 2004. Active oxygen species and antioxidants in seed biology. Seed Science Research14: 93–107.

Bailly C. 2019. The signalling role of ROS in the regulation of seed germination and dormancy.Biochemical Journal 476: 3019–3032.

Ballesteros D, Pritchard HW, Walters C. 2020. Dry architecture: towards the understanding of the variation of longevity in desiccation-tolerant germplasm. Seed Science Research 30: 142– 155.

Baskin CC, Baskin JM. 2014. Ecologically Meaningful Germination Studies In: Seeds. Elsevier, 5–35.

Begcy K, Mariano ED, Gentile A, et al. 2012. A Novel Stress-Induced Sugarcane Gene Confers Tolerance to Drought, Salt and Oxidative Stress in Transgenic Tobacco Plants (C Ng, Ed.). PLoS ONE 7: e44697.

Begcy K, Sandhu J, Walia H. 2018. Transient Heat Stress During Early Seed Development Primes Germination and Seedling Establishment in Rice. Frontiers in Plant Science 9: 1768.

Bewley JD, Bradford KJ, Hilhorst HWM, Nonogaki H. 2013. Development and Maturation In: Seeds. New York, NY: Springer New York, 27–83.

Capblancq T, Lachmuth S, Fitzpatrick MC, Keller SR. 2022. From common gardens to candidate genes: exploring local adaptation to climate in red spruce. New Phytologist: nph.18465.

Carta A, Bottega S, Spanò C. 2018. Aerobic environment ensures viability and anti-oxidant capacity when seeds are wet with negative effect when moist: implications for persistence in the soil. Seed Science Research 28: 16–23.

Chen D, Li Y, Fang T, Shi X, Chen X. 2016. Specific roles of tocopherols and tocotrienols in seed longevity and germination tolerance to abiotic stress in transgenic rice. Plant Science 244: 31– 39.

Cohen J. 1988. Statistical Power Analysis for the Behavioral Sciences. LEA Publishers.

Corbineau F. 2012. Markers of seed quality: from present to future. Seed Science Research 22: S61–S68.

Eckert AJ, Neale DB. 2022. Probing the dark matter of environmental associations yields novel insights into the architecture of adaptation. New Phytologist: nph.18639.

Ellis RH. 2019. Temporal patterns of seed quality development, decline, and timing of maximum quality during seed development and maturation. Seed Science Research 29: 135–142.

El-Maarouf-Bouteau H, Bailly C. 2008. Oxidative signaling in seed germination and dormancy. Plant Signaling & Behavior 3: 175–182.

Franks SJ, Richards CL, Gonzales E, Cousins JE, Hamrick JL. 2004. Multi-scale genetic analysis of Uniola paniculata (Poaceae): a coastal species with a linear, fragmented distribution. American Journal of Botany 91: 1345–1351.

Groot SPC, de Groot L, Kodde J, van Treuren R. 2015. Prolonging the longevity of *ex situ* conserved seeds by storage under anoxia. Plant Genetic Resources 13: 18–26.

Hacker SD, Jay KR, Cohn N, et al. 2019. Species-Specific Functional Morphology of Four US Atlantic Coast Dune Grasses: Biogeographic Implications for Dune Shape and Coastal Protection. Diversity 11: 82.

Halliwell B, Aeschbach R, Löliger J, Aruoma OI. 1995. The characterization of antioxidants. Food and Chemical Toxicology 33: 601–617.

Hart EM, Bell K. 2015. prism: Download data from the Oregon prism project.

Hodel RG, Gonzales E. 2013. Phylogeography of Sea Oats (Uniola paniculata), a Dune-building Coastal Grass in Southeastern North America. Journal of Heredity 104: 656–665.

Kim T, Samraj S, Jiménez J, Gómez C, Liu T, Begcy K. 2021. Genome-wide identification of heat shock factors and heat shock proteins in response to UV and high intensity light stress in lettuce. BMC Plant Biology 21: 185.

Lee Jae-Sung, Kwak J, Cho J-H, et al. 2019. A high proportion of beta-tocopherol in vitamin E is associated with poor seed longevity in rice produced under temperate conditions. Plant Genetic Resources: Characterization and Utilization 17: 375–378.

Lonard RI, Judd FW, Stalter R. 2011. Biological Flora of Coastal Dunes and Wetlands: Uniola paniculata L. Journal of Coastal Research 276: 984–993.

Long RL, Gorecki MJ, Renton M, et al. 2015. The ecophysiology of seed persistence: a mechanistic view of the journey to germination or demise: The ecophysiology of seed persistence. Biological Reviews 90: 31–59.

Malovichko YV, Shikov AE, Nizhnikov AA, Antonets KS. 2021. Temporal Control of Seed Development in Dicots: Molecular Bases, Ecological Impact and Possible Evolutionary Ramifications. International Journal of Molecular Sciences 22: 9252.

Martz F, Jaakola L, Julkunen-Tiitto R, Stark S. 2010. Phenolic Composition and Antioxidant Capacity of Bilberry (Vaccinium myrtillus) Leaves in Northern Europe Following Foliar Development and Along Environmental Gradients. Journal of Chemical Ecology 36: 1017–1028.

Moreira X, Abdala-Roberts L, Bruun HH, et al. 2020. Latitudinal variation in seed predation correlates with latitudinal variation in seed defensive and nutritional traits in a widespread oak species. Annals of Botany 125: 881–890.

Nguyen CD, Chen J, Clark D, Perez H, Huo H (Alfred). 2021. Effects of Maternal Environment on Seed Germination and Seedling Vigor of Petunia × hybrida under Different Abiotic Stresses. Plants 10: 581.

Oomah BD, Kenaschuk EO, Mazza G. 1997. Tocopherols in Flaxseed. Journal of Agricultural and Food Chemistry 45: 2076–2080.

Panuccio MR, Fazio A, Musarella AJ, Mendoza-fernández AF, Mota JF, Spampinato G. 2018. Seed germination and antioxidant pattern in Lavandual multifida (Lamiaceae): A comparison between core and peripheral populations. 152: 398–406.

Pehlivan FE. 2017. Free Radicals and Antioxidant System in Seed Biology In: Jimenez-Lopez JC, ed. Advances in Seed Biology. InTech,.

Pérez HE, Kane ME. 2017. Different plant provenance same seed tolerance to abiotic stress: implications for ex situ germplasm conservation of a widely distributed coastal dune grass (Uniola paniculata L.). Plant Growth Regulation 82: 123–137.

Peters, J., Lanham, B. 2000. Tetrazolium testing handbook, contribution No. 29 to the handbook on seed testing.

Re R, Pellegrini N, Proteggente A, Pannala A, Yang M, Rice-Evans C. 1999. Antioxidant activity applying an improved ABTS radical cation decolorization assay. Free Radical Biology and Medicine 26: 1231–1237.

Roach T, Nagel M, Börner A, Eberle C, Kranner I. 2018. Changes in tocochromanols and glutathione reveal differences in the mechanisms of seed ageing under seedbank conditions and controlled deterioration in barley. Environmental and Experimental Botany 156: 8–15.

Rogozhin VV, Verkhoturov VV, Kurilyuk TT. 2001. The Antioxidant System of Wheat Seeds during Germination. 28.

Ross KA, Zhang L, Arntfield SD. 2010. Understanding Water Uptake from the Induced Changes Occurred During Processing: Chemistry of Pinto and Navy Bean Seed Coats. International Journal of Food Properties 13: 631–647.

Saatkamp A, Cochrane A, Commander L, et al. 2019. A research agenda for seed-trait functional ecology. New Phytologist 221: 1764–1775.

Stegner M, Wagner J, Roach T. 2022. Antioxidant depletion during seed storage under ambient conditions. Seed Science Research: 1–7.

Subudhi PK, Parami NP, Harrison SA, Materne MD, Murphy JP, Nash D. 2005. An AFLP-based survey of genetic diversity among accessions of sea oats (Uniola paniculata, Poaceae) from the southeastern Atlantic and Gulf coast states of the United States. Theoretical and Applied Genetics 111: 1632–1641.

Vertucci CW, Farrant JM. 1995. Acquisition and Loss of Desiccation Torelance In: Kigel J, Galili G, eds. Seed Development and Germination. New York, NY: CRC Press Taylor & Francis Group, 237–271.

Walters C, Farrant JM, Pammenter NW, Berjak P. 2002. Desiccation Stress and Damage In: Black M, Pritchard HW, eds. Desiccation and survival in plants: drying without dying. Wallingford, Oxon, UK; New York: CABI Pub,.

Walters Christina, Hill LM, Wheeler LJ. 2005. Dying while Dry: Kinetics and Mechanisms of Deterioration in Desiccated Organisms. Integrative and Comparative Biology 45: 751–758.

Zeng Q, Dong G, Tian L, et al. 2020. High Altitude Is Beneficial for Antioxidant Components and Sweetness Accumulation of Rabbiteye Blueberry. Frontiers in Plant Science 11: 573531.

Zhou L, Yu H, Yang K, Chen L, Yin W, Ding J. 2021. Latitudinal and Longitudinal Trends of Seed Traits Indicate Adaptive Strategies of an Invasive Plant. Frontiers in Plant Science 12: 657813.

